# Gene regulatory mechanisms underlying evolutionary adaptations of homologous neuronal cell types

**DOI:** 10.1101/2025.01.30.635650

**Authors:** Andrea Millán-Trejo, Carlos Mora-Martínez, Adrián Tarazona-Sánchez, Carla Lloret-Fernández, Rafael Alis, Antonio Jordán, Arantza Barrios, Nuria Flames

## Abstract

How nervous systems coordinate the generation of specific neuron types with gene expression plasticity and how these mechanisms impact cell type evolution is unknown. Here we use *Caenorhabditis* species to study neuron-type robustness, plasticity and evolution, using VC4 and VC5 cholinergic motoneurons as models. In *C. elegans*, we found that epigenetic silencing through histone 3 lysine 9 methylation (H3K9me) is necessary to suppress the expression of the serotonin reuptake gene *mod-5/*Sert and a serotonergic phenotype in these cells. In contrast, we observed that VC4 and VC5 neurons in the *Angaria* group of species of the *Caenorhabditis* genus have evolved an intense serotonergic staining. This phenotype is caused by the emergence of a new enhancer in the *mod-5/*Sert locus, which has been recruited to the ancestral neuron-type gene regulatory network. Enhancer transfer from *C. angaria* is sufficient to impose a constitutive serotonergic fate in *C. elegans*. Remarkably, acquiring this new trait modulates egg-laying responses to high levels of exogenous serotonin, which can be found in specific environments. Finally, we discovered that the repression of the serotonergic fate in *C. elegans* VC4 and VC5 neurons is indeed a plastic trait that can be adjusted in specific environmental growth conditions to elicit egg-laying behaviours similar to those observed in *Angaria* species. Our work identifies gene regulatory mechanisms that coordinate the generation of robust neuron-type-specific programs with plastic gene expression responses. These findings identify a gene regulatory framework underlying the evolution of neuron-type-specific features and the emergence of novel behaviours.

---

Nervous systems are remarkably complex, containing an extraordinary array of cell types with highly specialised functions. The generation of this diversity is a deeply stereotyped process in which all individuals from the same species give rise to an identical variety of neuron types. Nonetheless, the evolution of nervous systems requires the emergence of novel neuronal identities, cellular functions, and behaviours. Changes in the architecture of gene regulatory networks have been proposed to play a key role in this evolutionary process^1,2^. However, the mechanisms controlling the robustness of neuronal development (i.e. the canalisation of phenotypes) and their impact on neuronal evolution have been challenging to delineate at the molecular level.

One example of neuronal diversity is found among serotonergic systems. In mammals, serotonin (5HT) is a widely distributed neurotransmitter controlling essential functions, including sensory processing, cognitive control, internal states, autonomic responses, and motor activity^3^. 5HT neurons can be classified as *bona fide* 5HT-synthesising if they contain the enzymes required for the synthesis of 5HT or 5HT-reuptaking if they express SERT, the transporter necessary to upload 5HT from the extracellular space^3^ (**Fig. 1a**). 5HT-synthesising neurons are restricted to the raphe region, where they are born during embryonic development and perdure in adulthood^3^. In contrast, 5HT-reuptaking neurons are found throughout the cortex, thalamus and retina only during development due to transient SERT expression in these neurons^3^. This ephemeral serotonergic feature has been involved in axon guidance, synapse formation or cell survival^3,4^. However, mechanisms controlling the temporally-restricted 5HT reuptake phenotype are not well understood^5^.

**Figure 1.**
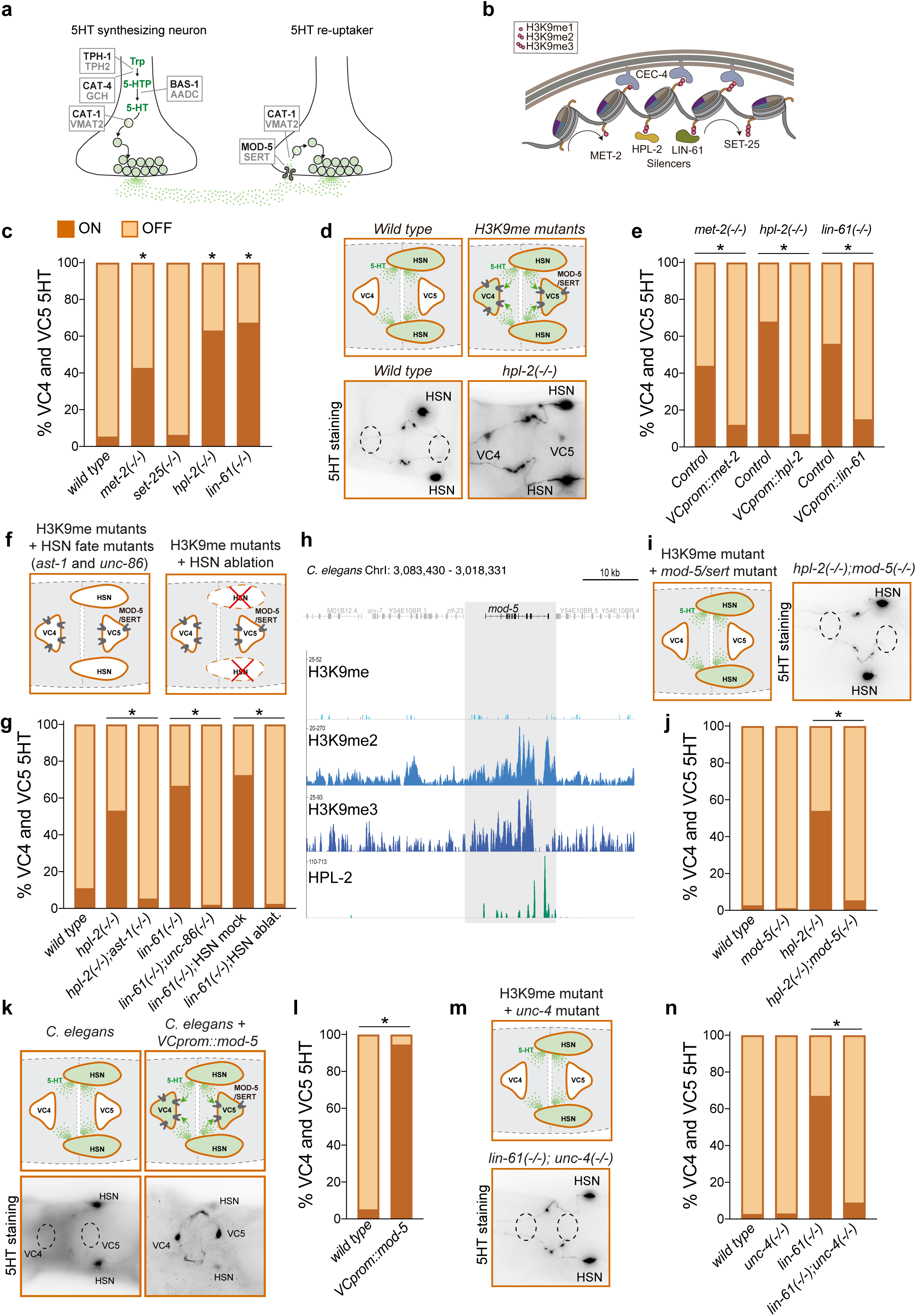
Epigenetic silencing prevents VC4 and VC5 serotonergic phenotype in *C. elegans*. **a)** Schema of the minimal set of proteins required for the synthesis (left) or reuptake and package (right) of 5HT. Black: *C. elegans* names, Grey: Human names. b) Schematic representation of main components of Histone 3 lysine 9 methylation pathway in *C. elegans*. **c**) VC4 and VC5 5HT staining quantification for H3K9me pathway mutants *met-2(n4256), set-25(n5021*), *lin-61(n3809*) and *hpl-2(tm1489).* Fisher’s exact test was applied for statistics. **d**) Schematic representation and representative micrographs of VC4 and VC5 5HT staining in *wild type* and H3K9me mutants. **e**) Quantification of H3K9me pathway mutants rescue experiments using VC specific promoter shows epigenetic repression of serotonergic phenotype is cell autonomous. Fisher’s exact test was applied for statistics. **f**) Schematic representation of experimental approaches used to eliminate 5HT from HSN neurons in H3K9me mutants. **g)** Quantification of VC4 and VC5 serotonergic phenotype in wild type, H3K9me mutants *hpl-2* and *lin-61* and double mutants *hpl-2(tm1489); ast-1(ot417)*, *lin-61(n3809); unc-86(n846)* and *lin-61(n3809*) with HSN laser ablation. **h***) C. elegans* larval stage L4 ChIPseq mono-, di- and tri-methylation levels and HPL-2 binding. *mod-5* locus (marked with grey shadow) shows high me1, me2, me3 and HPL-2 recruitment. **i)** Schematic representation and micrograph of VC4 and VC5 5HT staining in double mutant H3K9me pathway and *mod-5(knu383)**. j)*** Quantification of (i), VC4 and VC5 serotonergic phenotype in H3K9me mutants requires *mod-5* activity. Fisher’s exact test was applied for statistics. **k**) Schematic depiction and representative micrographs of 5HT staining in C*. elegans* control animals and animals expressing *mod-5/*Sert cDNA under a VC specific promoter. Expression of *mod-5/*Sert is sufficient to drive serotonergic phenotype in *C. elegans. **l)*** Quantification of (k). Fisher’s exact test was applied for statistics**. m)** Schematic depiction and representative micrographs of 5HT staining of double mutant H3K9me pathway and *unc-4(gk705*). **o**) Quantification of (m), Fisher’s exact test was applied for statistics. See Supplementary information for raw data and p values.

In animals, the use of 5HT as a neurotransmitter dates to the bilaterian origin, present in annelids, molluscs, arthropods, nematodes, and vertebrates^6^. Assignments of neuronal homologies across species are required to study neuron-type evolution. Nevertheless, this process can be challenging in complex organisms due to significant anatomical differences. Advantageously, all nematode species in the *Caenorhabditis* genus share a common invariable embryonic lineage that gives rise to a strikingly identical number of neurons placed in the same body locations^7,8^. Thus, the genetic amenability of *C. elegans* and the existence of closely related *Caenorhabditis* nematodes offer a unique opportunity to investigate the generation of cell diversity and the emergence of novel neuronal features underlying key evolutionary steps. *C. elegans* contains multiple serotonergic neuron types that regulate sensory processing, motor function and internal states^9,10^. As in mammals, these cells can be classified as 5HT-synthesizing and 5HT-reuptaking neurons, although, in contrast to mammals, *C. elegans* 5HT reuptake neurons are maintained throughout the life of the animal^11,12^. Two cholinergic motoneurons in *C. elegans* (VC4 and VC5 neuron types) have been reported to display weak serotonin staining^13,14^. However, this staining was described as sporadic and variable among individuals and experiments, contrasting with the stereotypical and robust classification of all other neuron types in *C. elegans*^15^. This discrepancy prompted us to investigate the molecular mechanisms underlying the sporadic 5HT staining in VC4 and VC5 neurons and the evolvability of this phenotype.

Here, we found that active epigenetic silencing is employed by *C. elegans* VC4 and VC5 neurons to prevent a 5HT reuptake phenotype. In contrast, VC4 and VC5 neurons in *Caenorhabditis* species of the *Angaria* group exhibit intense 5HT staining due to the emergence of a new enhancer driving constitutive *mod-5/Sert* expression in these cells. We also unravelled that the VC4 and VC5 5HT phenotype explains specific differences in egg-laying behaviour between *C. elegans* and *C. angaria* species. Finally, we revealed that the repression of the VC4 and VC5 serotonergic phenotype in *C. elegans* is a plastic trait that can be modulated, and we identified the specific environmental conditions that overcome epigenetic silencing of *mod-5/Sert* in this species, resulting in egg-laying behaviours like those found in *C. angaria*. In sum, our results highlight the importance of epigenetic repression in the canalisation of neuron-type identities, illuminate specific gene regulatory changes driving the evolution of novel neuronal phenotypes and behaviours, and identify unique mechanisms of neuronal identity plasticity in response to environmental signals. These findings illustrate how the modulation of repressive mechanisms impacts developmental plasticity and the evolution of nervous systems.

## Epigenetic silencing restricts VC4 and VC5 serotonergic phenotype in C. elegans

As previously reported^13,14^, weak 5HT staining in VC4 and VC5 neurons was observed in a low percentage of *C. elegans* N2 animals under our standard laboratory conditions (**Fig. 1c**), in contrast to robust staining found in all known 5HT synthesising (ADF, NSM and HSN) and reuptaking (AIM and RIH) neurons in *C. elegans*. The epigenetic mark Histone H3 lysine 9 methylation (H3K9me) is associated with heterochromatin and transcriptional silencing^16^ (**Fig. 1b**). Recent reports indicate that H3K9me can be deposited in a cell-type-specific manner to repress expression of undesired effector features of related cellular lineages, like olfactory receptor repression in the diversification of olfactory receptor neurons^16–21^. We find that mutants for different H3K9me pathway components^22^ (specifically alleles for mono- and dimethyl transferase *met-2/*Setdb1 and *hpl-2/*Hp1 and *lin-61/*L3mbtl2 protein silencers) show a strong, reproducible and highly penetrant 5HT staining in VC4 and VC5 neurons (**Fig. 1c, d and Source Data**). The VC4 and VC5 serotonergic phenotype is rescued with the re-expression of the corresponding gene under a VC neuron-specific promoter supporting a cell-autonomous role for H3K9me (**Fig. 1e and Source Data**). Next, we aimed to determine if VC4 and VC5 become 5HT synthesising or reuptaking neurons in these mutant backgrounds. Wildtype *C. elegans* VC4 and VC5 neurons constitutively express the vesicular monoamine transporter *cat-1/*Vmat^13^. Of note, the function of this gene in the otherwise cholinergic VC4 and VC5 motoneurons remains undetermined. Accordingly, the acquisition of *mod-5/*Sert 5HT expression should be sufficient to acquire the serotonergic phenotype (**Fig. 1a**). Thus, we considered 5HT reuptake from nearby HSN 5HT synthesising neurons the more plausible scenario. To test this hypothesis, we built double mutants for H3K9me pathway components together with mutants for transcription factors necessary for HSN differentiation, which are not expressed in VC4 and VC5 neurons (*ast-1/*Fli1^23^ and *unc-86/*Brn3c^24,25^). We find that these genetic backgrounds display a complete absence of VC4 and VC5 5HT staining (**Fig. 1f, g** and **Source data**). In complementary experiments, we used laser ablation to eliminate HSN neurons in *lin-61/*L3mbtl2 mutant animals before HSN maturation at the first larval stage. HSN depletion leads to substantial defects in VC4 and VC5 5HT staining (**Fig. 1f, g** and **Source data**), further supporting a reuptake phenotype for these neuron types in H3K9me mutants. Interrogation of available chromatin immunoprecipitation and sequencing (ChIP-seq) data^26^ reveals *C.elegans* wildtype animals show high H3K9me marks and HPL-2/HP1 recruitment to the *mod-5/*Sert locus (**Fig. 1h**). Double mutant analysis of *hpl-2/*Hp1 with *mod-5/*Sert alleles confirms VC4 and VC5 neurons reuptake 5HT in a *mod-5/*Sert dependent manner (**Fig. 1i, j and Source data**). However, we failed to detect *mod-5/*Sert expression in VC4 and VC5 neurons in H3K9me pathway mutants using an endogenously tagged *mod-5::T2A::NeonGreen* strain (**Extended Data Fig. 1**), which could be due to low levels of expression. Nevertheless, we found that induced *mod-5/*Sert expression in otherwise wildtype VC4 and VC5 neurons is sufficient to drive intense 5HT staining (**Fig. 1k, l** and **Source data**). Altogether, our results support a model in which *mod-5/*Sert expression in VC4 and VC5 neurons is actively epigenetically repressed by the H3K9me pathway, suppressing the 5HT reuptake phenotype.

## Low UNC-4/UNCX restricts VC4 and VC5 serotonergic phenotype in *C. elegans*

Epigenetic modifiers act in concert with transcription factors (TFs) to activate or repress specific target genes^17^. Thus, we next aimed to identify TFs involved in activating the VC4 and VC5 5HT reuptake phenotype in H3K9me pathway mutants. The homeodomain TF UNC-4, a homolog of vertebrate UNCX, is required for the terminal differentiation of VC4 and VC5 cholinergic neurons ^27–29^. We found that UNC-4/UNCX is also necessary for constitutive *cat-1/*Vmat reporter expression in VC4 and VC5 neurons (**Extended Data Fig. 1**). Analysis of the *cat-1/*Vmat minimal *cis* regulatory module (CRM or enhancer) active in VC4 and VC5 neurons revealed the presence of a functional homeodomain TF binding site, suggesting UNC-4/UNCX action could be direct (**Extended Data Fig. 1**). In addition, VC4 and VC5 5HT staining is absent in *unc-4/*Uncx*; lin-61/*L3mbtl2 double mutants, revealing UNC-4/UNCX is necessary for serotonergic phenotype induction in H3K9me mutants (**Fig. 1m, n and Source data**). Considering these results, we next aimed to assess if increased UNC-4/UNCX levels could overcome H3K9me repression. We observed that UNC-4/UNCX overexpression through a heat shock inducible promoter is sufficient to induce 5HT staining in *C. elegans* VC4 and VC5 neurons (**Extended Data Fig. 1**). Similar to H3K9me pathway mutants, we failed to detect VC4 and VC5 *mod-5/*Sert expression using endogenously tagged *mod-5::T2A::NeonGreen* strain (**Extended Data Fig. 1**). However, two pieces of evidence suggest UNC-4/UNCX overexpression acts through *mod-5/*Sert induction: first UNC-4/UNCX overexpression induced serotonergic phenotype is absent in *mod-5/*Sert mutants (**Extended Data Fig. 1**), and second, high levels of UNC-4/UNCX are sufficient to induce detectable VC4 and VC5 GFP expression from the multicopy array *mod-5prom3::gfp* transgenic reporter which, under control conditions, is only express in NSM and ADF serotonergic synthesizer neurons (**Extended Data Fig. 1**). Finally, we examined if serotonergic phenotype of H3K9me mutants is mediated through increasing UNC-4/UNCX levels, however quantification of an endogenous fluorescence tagged *unc-4/*Uncx allele reveals a slight but significant decreased in UNC-4/UNCX levels (**Extended Data Fig. 1**). This observation is consistent with previous data showing that H3K9me restricts binding of TF fate determinants to unwanted target effector genes but does not affect TF expression^17^. Altogether, our results support a model in which *C. elegans mod-5/*Sert locus bares some latent enhancer activity in VC4 and VC5 neurons that is actively prevented by two parallel mechanisms, restricted UNC-4/UNCX expression levels and epigenetic H3K9me repression.

## VC4 and VC5 serotonergic phenotype in *Angaria* group of species

The serotonergic phenotype of VC4 and VC5 neurons has been reported in other nematode species^30^ but has not been systematically studied in the *Caenorhabditis* genus. We performed 5HT immuno-staining in 26 *Caenorhabditis* species (**Fig. 2a, b**) and found highly penetrant and strong VC4 and VC5 5HT staining in *C. angaria*, as previously reported^30^, and in four additional species of the *Angaria* group (**Fig. 2a, b** and **Source data**). These data suggest that VC4 and VC5 neurons acquired the ability to contain 5HT in the common ancestor of the *Angaria* group (synapomorphy). Next, we aimed to identify the molecular determinants of this acquired trait.

**Figure 2.**
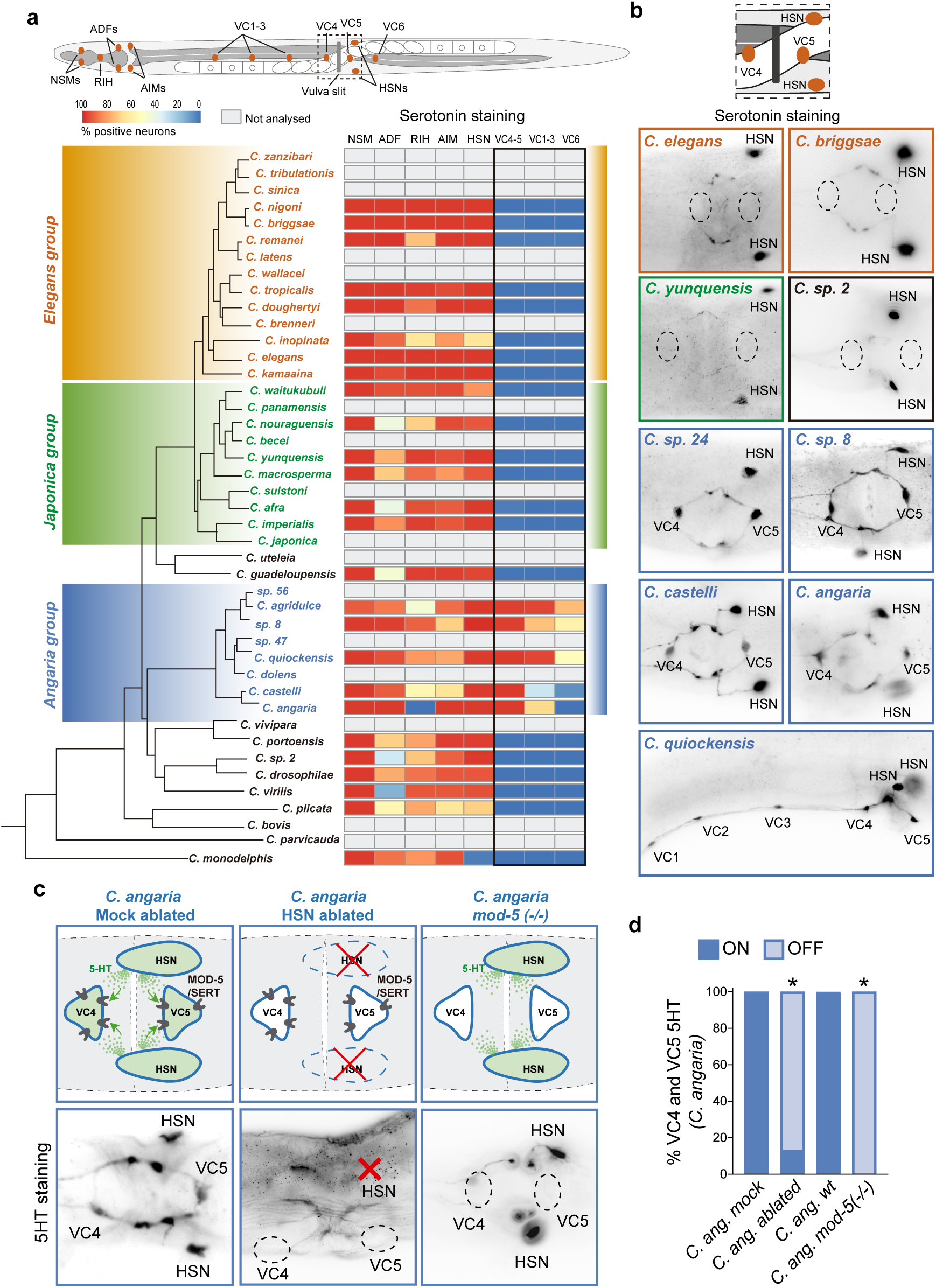
VC4 and VC5 neurons from *Angaria* group of species are *mod-5/*Sert dependent serotonin reuptakers. **a)** Schema of soma locations for serotonergic and VC neurons in *Caenorhabditis* species and phylogeny of *Caenorhabditis* species (adapted from http://caenorhabditis.org/). Heatmap quantification of serotonergic staining. VC4 and VC5 neurons display strong 5HT staining in all analysed *Angaria* group of species. n>25 animals per specie. **b)** Micrographs showing illustrative examples of 5HT staining in the vulva region of several species. **c)** Schematic depiction (top) and representative micrographs (bottom) of 5HT staining in C*. angaria* mock ablated animals, HSN ablated animals and *C.angaria mod-5(vcl45)* mutants. Loss of HSN or *mod-5* gene abolish VC4 and VC5 5HT staining**. d)** quantification of experiments in (c). Fisher’s exact test was applied for statistics. See Supplementary information for raw data and p values.

## *Angaria* VC4 and VC5 neurons reuptake serotonin from HSN neurons

To determine if, like in *C. elegans* H3K9me mutants, the serotonergic phenotype of *C. angaria* VC4 and VC5 is due to 5HT reuptake, we eliminated the HSN neurons by laser ablation. *C. angaria* HSN ablated animals show reduced VC4 and VC5 5HT staining compared to mock ablated controls (**Fig. 2c, d** and **Source data**). We next generated *C. angaria mod-5/*Sert null animals *[Cang-mod-5(vlc45)]* (**Extended Data Fig. 2**). We found a complete absence of 5HT labelling in VC4 and VC5 neurons in these mutants, which reinforced the view that these cells reuptake 5HT produced by HSN neurons (**Fig. 2c, d**). Like in *C. elegans*, *C. angaria* VC4 and VC5 neurons stain for the UNC-17/VCHAT vesicular acetylcholine transporter, suggesting that they maintain their cholinergic identity while acquiring a serotonergic phenotype rather than undergoing a fate switch (**Extended Data Fig. 2**). Dual neurotransmission is present in other *C. elegans* neuron types^13^. Altogether, these experiments imply that VC4 and VC5 neurons in the *Angaria* group of species have acquired robust *mod-5*/Sert expression, which allows them to reuptake 5HT from nearby HSN neurons.

## Regulatory regions of *C. angaria mod-5*/Sert are active in *C. angaria* and *C. elegans* VC4 and VC5 neurons

We next aimed to identify the molecular mechanisms driving the expression of *mod-5/*Sert in VC4 and VC5 neurons in *Angaria* group of species. Because *C. angaria* is not a model organism and we failed to systematically generate stable transgenic lines, alternatively we carried out the analysis in F1 transgenic animals. As expected, we found regulatory regions of *C. angaria mod-5/*Sert locus (*Cang_mod-5* reporter) active in *C. angaria* VC4 and VC5 neurons, supporting a cell autonomous role for *mod-5/*Sert in the acquisition of the serotonergic phenotype (**Fig. 3c, Source data** and **Extended Data Fig. 3**).

**Figure 3.**
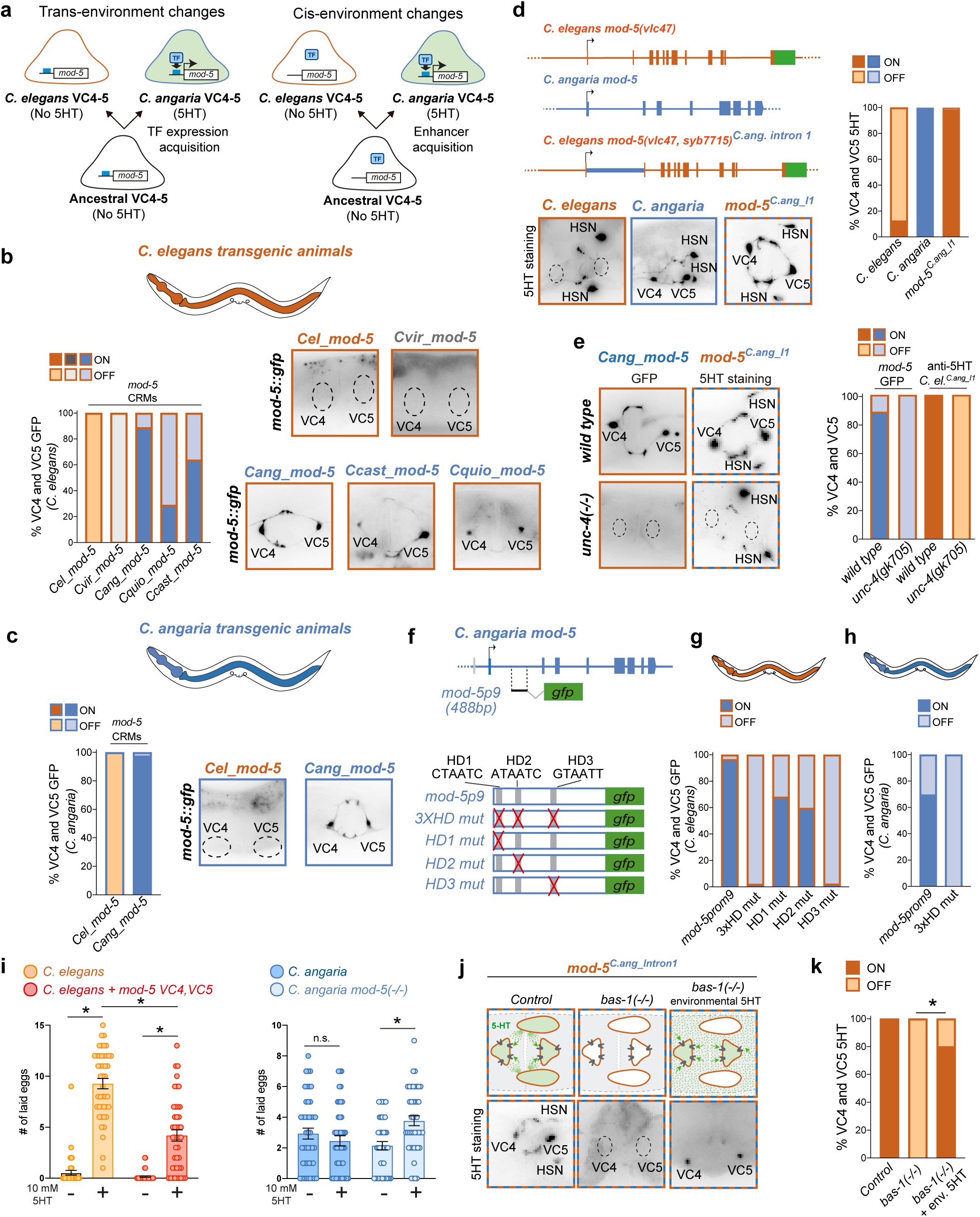
cis regulatory changes underlie VC4 and VC5 serotonergic phenotype in *Angaria* species and provide differential egg laying responses. **a)** Schematics of two models explaining the emergence of *mod-5/*Sert expression in *Angaria* group of species. **b**) *C. elegans* transgenic reporter animals are used to study *mod-5/*Sert regulatory region activity from different *Caenorhabditis* species: two non-angaria species *C. elegans (Cel)* and *C.virilis (Cvir)* and three *Angaria* species *C. angaria (Cang), C. quickensis (Cquio)* and *C. castelli (Ccast). mod-5/*Sert CRMs from *Angaria* species but not from other species drive *gfp* expression in *C. elegans* VC4 and VC5 neurons. Graph represents the mean of at least 2 lines n>60. c) *C. angaria* F1 transgenic reporter animals are used to study *C. elegans* and *C. angaria mod-5/*Sert regulatory region activity*. C. angaria* but not *C. elegans* reporter is active in *C. angaria* VC4 and VC5 neurons n=33;28 respectively. **d)** Schematic representation of *mod-5/*Sert locus in *C. elegans* and *C. angaria. Cel mod-5(vlc47)* corresponds to a CRISPR tagged *T2A::NeonGreen* allele, while *Cel mod-5(syb7715)* substitutes *Cel* intron1 sequence by *Cang* intron1. Micrographs and quantification of 5HT staining shows that intron 1 swap is sufficient to drive strong 5HT phenotype in *C. elegans* VC4 and VC5 neurons. **e)** Micrographs and quantification of *Cang_mod-5*::*gfp* reporter expression and 5HT staining of *Cel mod-5(vlc47,syb7715)* in *wild type* and *unc-4(gk705)* mutant background. UNC-4 is required for VC4 and VC5 serotonergic fate. (f) *C. angaria mod-5* locus schema and location of the minimal CRM (*mod-5prom9*) driving expression in VC4 and VC5 neurons. *mod-5prom9* contains three putative HD binding sites. Individual and combined HD point mutations were tested for its activity in *C. elegans* and *C. angaria* transgenic animals. **g)** Quantification of mutated constructs in *C. elegans* reporter lines. **h)** Quantification of mutated constructs in *C. angaria* transgenic animals**. i**) Quantification of egg laying events upon 2h exposure to external 5HT. Both, *C. elegans* control animals (N2) and transgenic animals with *mod-5* cDNA expression in VC neurons induce egg laying upon 5HT exposure, however, egg laying induction in *mod-5* transgenic animals is significantly lower to controls*. C. angaria* control animals (RGD1) do not increase egg laying when exposed to external 5HT while *C. angaria mod-5* mutant animals show significant egg laying induction. Nonparametric Mann–Whitney test. **j**) Schematic depiction (top) and representative micrographs (bottom) of VC4 and VC5 neuron 5HT staining in *Cel mod-5(vlc47,syb7715)* animals, *Cel mod-5(vlc47,syb7715); bas-1(ad446)* animals that lack 5HT biosynthesis and *Cel mod-5(vlc47,syb7715); bas-1(ad446)* animals exposed to high levels of exogenous 5HT**. k**) Quantification of (j) shows VC4 and VC5 neurons can uptake 5HT from exogenous sources. Fisher’s exact test. See supplementary Information for raw data and p values.

Two alternative scenarios could explain mod*-5/*Sert expression in VC4 and VC5 neurons in *C. angaria*. First, changes in the trans-environment (e.g., differential expression of TFs or epigenetic landscape in VC4 and VC5 neurons in *C. angaria* versus *C. elegans*) could be responsible for *de novo* activation of *mod-5/*Sert gene (**Fig. 3a, left**). Second, changes in the cis-regulatory regions of *mod-5/*Sert locus might result in the emergence of a new enhancer that resulted in its activation by the ancestral VC4 and VC5 trans-environment (**Fig. 3a, right**). These possible scenarios make specific predictions for the outcome of heterologous gene reporter experiments. If trans-environment changes are mainly responsible for the phenotype, *C. elegans* regulatory region reporters (*Cel_mod-5/*Sert) would be activated in *C. angaria* VC4 and VC5 neurons while *C. angaria* reporter (*Cang_mod-5/*Sert) would be inactive in the context of *C. elegans* neurons. Conversely, if *cis*-regulatory changes are the main drivers of the phenotype, the *Cang_mod-5/*Sert reporter would be active in *C. elegans* VC4 and VC5 neurons and *Cel_mod-5/*Sert would not be active in *C. angaria*. Our heterologous reporter analyses supported this second scenario: a cis-environment model, as species origin of the regulatory regions, dictates expression (*Cang_mod-5/*Sert ON; *Cel_mod-5/*Sert OFF) and not the species-specific VC4 and VC5 cellular context (**Fig. 3b, c, Extended data Fig. 3** and **Source data**). Further supporting the cis-model, *mod-5/*Sert reporters from two additional *Angaria* species (*C. castelli* and *C. quiockensis*) show VC4 and VC5 activation in *C. elegans* transgenic animals, like the *C. angaria* construct, while an equivalent reporter from *C. virilis,* an *Angaria* outgroup specie, does not (**Fig. 3b, Extended data Fig. 3 and Source data**). In summary, our results show that *Angaria* species share the presence of a CRM in the *mod-5/*Sert locus, which is constitutively activated in VC4 and VC5 neurons by transcription factor/s shared between *Angaria* species and *C. elegans*.

## *C. angaria mod-5/*Sert enhancer is sufficient to provide serotonergic phenotype to *C. elegans* VC4 and VC5 neurons

*C. angaria mod-5/*Sert CRM active in VC4 and VC5 neurons is in the first intron of the locus (**Extended Data Fig. 3**). In *C. elegans*, the *Cel_mod-5 intron1* reporter is active in NSM and ADF but not VC4 and VC5, while the *Cang_mod-5 intron1* reporter is active in NSM, VC4 and VC5 (**Extended Data Fig. 3 and Source data**). To test sufficiency of the *C. angaria* enhancer in its endogenous genomic context we modified the *C. elegans mod-5::T2A::NeonGreen* strain *[mod-5(vlc47)]* substituting the first intron by *C. angaria* intron 1 sequence [generating *C. elegans mod-5(vlc47syb7715)* referred as *Cel*_*mod-5^C.ang_intron^*^1^] (**Fig. 3d**). This intron swap is sufficient to drive the VC4 and VC5 serotonergic phenotype in *C. elegans* like in *C. angaria* (**Fig. 3d**), further supporting the *cis*-regulatory model. These results suggested that the ancestral *Angaria* species gained robust *mod-5/*Sert expression in VC4 and VC5 neurons through the emergence of an enhancer that overcomes H3K9me repression, allowing its recruitment to the pre-existing VC4 and VC5 gene regulatory network.

## UNC-4/UNCX directly activates *C. angaria mod-5/*Sert enhancer

Next, we aimed to identify TFs responsible for activating the *C. angaria mod-5/*Sert enhancer. Since genetic manipulations in *C. angaria* are challenging and the *Cang_mod-5 intron 1* enhancer is active in VC4 and VC5 neurons in both *C. angaria* and *C. elegans*, we hypothesised that the TFs responsible for its activation are conserved between both species. Based on this idea, we used *C. elegans* for the initial characterisation of its regulatory logic. Consistent with our findings in *C. elegans,* where UNC-4/UNCX is necessary for *C. elegans cat-1::gfp* expression in VC4 and VC5 neurons and the serotonergic phenotype in H3K9me mutants, we observed that the *Cang_mod-5::gfp* fluorescence reporter is inactive in VC4 and VC5 neurons in *C. elegans unc-4/*Uncx*(gk705)* mutants, while reporter expression in NSM neurons is unaffected (**Fig. 3e** and **Source data**). Consistently, *C. elegans mod-5^C.ang_intron^*^1^ strain loses VC4 and VC5 5HT staining in *unc-4/*uncx*(gk705)* mutant background (**Fig. 3e**). UNC-4/UNCX activation of *C. angaria mod-5*/Sert enhancer is likely direct because point mutations in predicted UNC-4/UNCX homeodomain binding sites (TAAT) abolished *Cang_mod-5* reporter gene expression both in *C. elegans* and *C. angaria* transgenic animals (**Fig. 3f, g, h** and **Source data**). These results suggest that *cis*-regulatory changes in *mod-5/*Sert intron 1 allowed the recruitment of this gene in *Angaria* as a new robust direct target of UNC-4/UNCX, which provided a constitutive VC4 and VC5 serotonergic phenotype in these species.

## The VC4 and VC5 serotonergic phenotype in *Angaria* species buffers egg-laying induction by environmental serotonin

Next, we aimed to identify behavioural differences that could be attributed to the acquisition of the serotonergic phenotype by VC4 and VC5 neurons. In *C. elegans,* VC4, VC5 and HSN motoneurons are part of the egg-laying circuit^31^. We found that, under laboratory conditions, the egg-laying behaviour of *C. elegans* and *C. angaria* strains is significantly different (**Extended Data Fig. 4**). However, many factors could explain these differences and not just those related to the serotonergic phenotype of VC4 and VC5 neurons. Indeed, when we specifically tested the contribution of *mod-5*/Sert expression in VC4 and VC5, we found egg laying is similar in *C. elegans* between wildtype and transgenic animals expressing *mod-5/*Sert in VC4 and VC5 neurons. Similarly, no apparent differences were observed in *C. angaria* between wild type and *mod-5/*Sert*(vlc45)* mutants (**Extended Data Fig. 4**).

We reasoned that laboratory growth conditions, which vastly differ from those of the nematode’s natural habitat, could explain the absence of a differential phenotype. *Caenorhabditis* species have been isolated mostly from rotten fruits and flowers, where they are exposed to complex chemical and microbial environments^32^. 5HT is found across all domains of life, including plants, bacteria and fungi, and it is consistently implicated in interspecies crosstalk, including plant defence against insects^33–39^. Considering that biotic interactions are often significant drivers of evolutionary change^40^, we hypothesised that *C. angaria* may encounter habitats with high levels of environmental 5HT and wondered whether this feature might be related to the acquired VC4 and VC5 serotonergic phenotype. Indeed, some *Angaria* species have been isolated from fruits with high 5HT content, such as tomatoes (*C. sp.8*) or associated with the weevil pests of bananas, which are also high in 5HT^41,42^. It is also known that the sensory behaviour of *C. elegans* can be modulated by bacterially synthesised neurotransmitters^43^, and in the laboratory, *C. elegans* exposure to exogenous 5HT induces egg laying^44^. Thus, to test this hypothesis, we first compared egg-laying responses to acute treatment with exogenous 5HT between wildtype *C. elegans* and *C. elegans* animals expressing *mod-5/*Sert in VC4 and VC5 neurons. Our results revealed that *C. elegans* egg-laying induction by exogenous 5HT is significantly attenuated by *mod-5/*Sert expression in VC4 and VC5 neurons (**Fig. 3i** and **Source data**). Conversely, *C. angaria* wildtype animals are resistant to egg-laying induction by exogenous 5HT while *C. angaria mod-5/*Sert mutants lose this buffering capacity (**Fig. 3i**). An extended analysis of egg-laying responses to exogenous 5HT in different *Caenorhabditis* species shows a strong anti-correlation between VC4 and VC5 5HT staining and egg-laying induction by exogenous 5HT (**Extended Data Fig. 5**). Altogether, our results suggested that the ability of VC4 and VC5 neurons to reuptake 5HT can prevent egg-laying induction by exogenous 5HT.

## *mod-5/*Sert expression in VC4 and VC5 neurons in *C. elegans* allows for environmental 5HT reuptake

How environmental 5HT induces acute egg laying in *C. elegans* is not known. In *C. elegans* 5HT released from HSN neurons stimulates egg laying by directly modulating the functional state of the vulval muscles^45–47^. We hypothesise that exogenous 5HT could reach vulval muscle serotonergic synapses either by diffusion through the vulva slit or ingestion, inducing vulval muscle contraction and egg-laying behaviour. We reasoned that in *Angaria* species, VC4 and VC5 neurons could not only reuptake 5HT from HSN synapses but also, if present, from external sources, thereby preventing excessive activation of neuromuscular synapses. To test this hypothesis, we took advantage of *C. elegans mod-5^C.ang_intron^*^1^ animals, which show intense VC4 and VC5 5HT staining under laboratory conditions, like *C. angaria*, and generated a double mutant with the *bas-1/*Aadc*(ad446)* allele*. bas-1/*Aadc codes for the amino acid decarboxylase enzyme, fundamental for endogenous 5HT biosynthesis^48^. As expected, under standard laboratory conditions *C. elegans mod-5^C.ang_intron^*^1^; *bas-1/*Aadc*(ad446)* animals show a complete loss of 5HT staining, including VC4 and VC5 neurons. However, these animals recover strong VC4 and VC5 5HT staining when exposed to exogenous 5HT (**Fig. 3j, k**), demonstrating that these neurons can uptake environmental 5HT. Together with our previous results, these observations suggested that the acquisition of *mod-5/*Sert expression in VC4 and VC5 neurons by the *Angaria* group of species had modified their egg-laying behaviour, allowing them to buffer egg-laying induction produced by exogenous 5HT.

## Environmental serotonin induces VC4 and VC5 serotonergic phenotype in *C. elegans*

*C. elegans* nervous system can adapt to different environmental stimuli through neuron-type specific changes in gene expression^15^. Considering the active epigenetic repression of *mod-5/*Sert locus in VC4 and VC5 neurons, we wondered if the serotonergic phenotype in *C. elegans* could represent a plastic trait and aimed to identify potential environmental triggers. As we determined that this trait in Angaria species helps prevent egg-laying induction by exogenous 5HT, we wondered if this environment could precisely trigger the VC4 and VC5 5HT phenotype in *C. elegans*. We found that *C. elegans* animals grown for one generation in high levels of environmental 5HT show a significant increase in VC4 and VC5 serotonergic staining (**Fig. 4a-c** and **Source data**). Induction of *C. elegans* VC4 and VC5 5HT staining requires UNC-4/UNCX and MOD-5/SERT, like in *C. elegans* H3K9me mutants and wildtype *C. angaria* (**Fig. 4b, c and Extended Data Fig. 6**). To assess how broadly distributed in the *Caenorhabditis* genus this plastic trait is, we exposed two additional species, *C. briggsae* and *C. tropicalis,* to high 5HT environments for one generation. We failed to detect VC4 and VC5 5HT staining in these animals, suggesting VC4 and VC5 plasticity to environmental 5HT is not a general feature of the *Caenorhabditis* genus (**Extended Data Fig. 6**). In addition, in contrast to *C. elegans*, *cat-1*/Vmat expression in VC4 and VC5 neurons is not detected in *C. briggsae* and *C. tropicalis*^49^, consistent with the lack of serotonergic phenotype plasticity.

**Figure 4.**
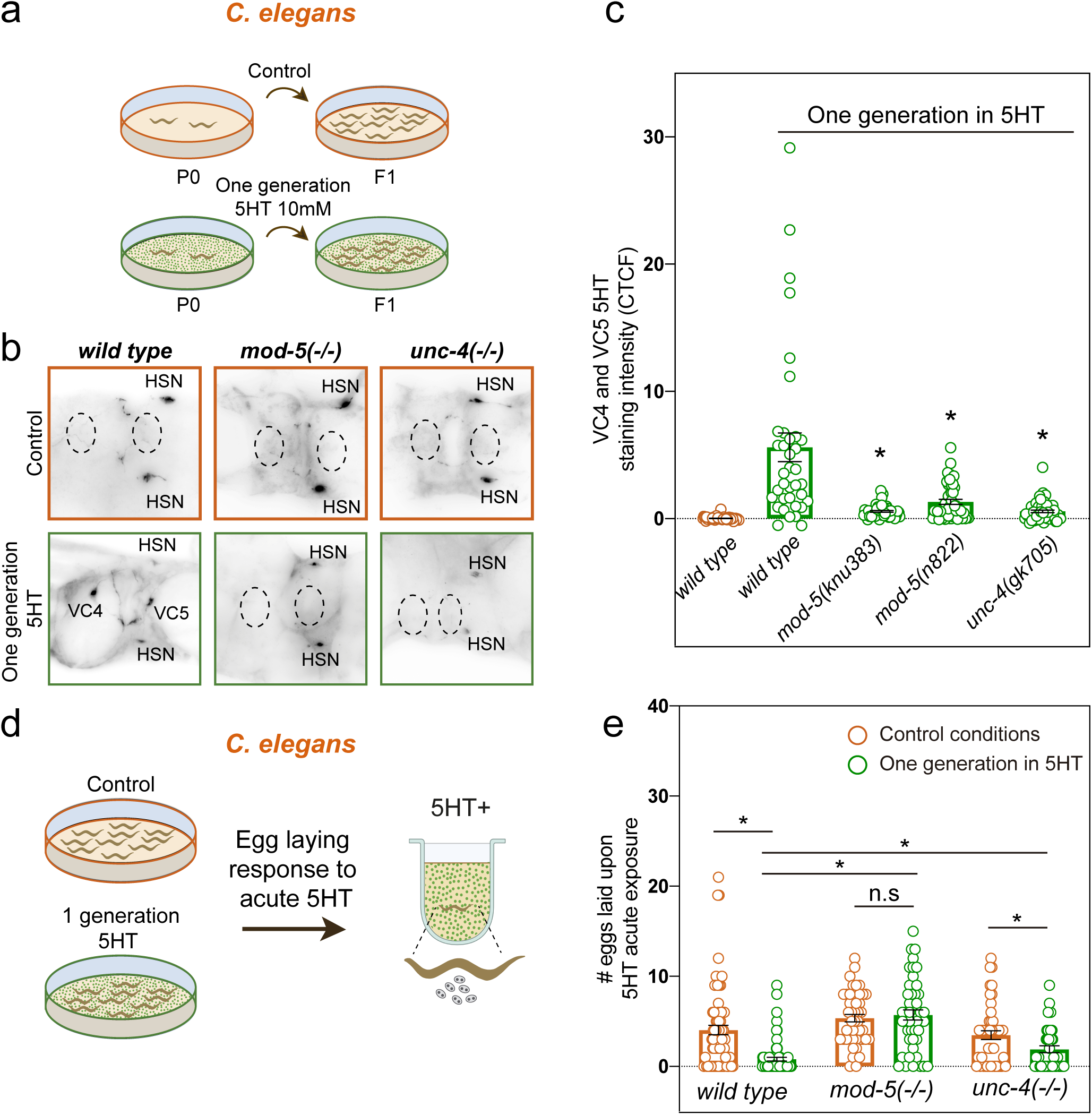
*C. elegans* VC4 and VC5 serotonergic phenotype is a plastic trait released by environmental 5HT exposure that leads to modified egg laying responses. **a)** Schematic representation of the experimental set up, worms are grown for one generation in control conditions or exposed to high levels of environmental 5HT. **b)** Representative micrographs of VC4 and VC5 serotonergic staining in *wild type*, *mod-5(knu383)* or *unc-4(gk705*) animals under control conditions or grown for one generation in environmental 5HT. **c)** Quantification of VC4 and VC5 serotonergic intensity phenotype. 5HT staining induction by environmental 5HT is significantly reduced in two different alleles of *mod-5/*Sert transporter and *unc-4(gk705)* mutants. Kustal-Wallis nonparametric test with Dunn’s correction for multiple comparisons. **d)** Schematic representation of the experimental set up, worms grown for one generation in control conditions or environmental 5HT are tested for acute egg laying responses (2h) to exogenous 5HT. **e)** Quantification of laid eggs. Wild type worms grown for one generation exposed to environmental 5HT (green) are able to buffer the egg laying induction upon acute 5HT exposure compared to controls (orange). This buffering effect is absent in *mod-5(knu383)* and reduced in *unc-4(gk705*) mutant animals. Mann-Whitney test. CTCF: Corrected Cell Total Fluorescence. See supplementary Information for raw data and p values.

Finally, we tested the behavioural consequences of this form of plasticity in *C. elegans*. We found that *C. elegans* animals grown in high 5HT environments modified their egg-laying responses to acute 5HT treatment in a way analogous to the constitutive behaviour observed in *Angaria* species, i.e., they buffer egg-laying induction by exogenous 5HT (**Fig 4d, e** and **Source data**). Importantly, this attenuation response is absent in *mod-5/*Sert mutants and significantly reduced in *unc-4/*Uncx mutant animals grown under similar 5HT exposure conditions, further supporting the requirement of *mod-5/*Sert expression in VC4 and VC5 neurons to mediate this form of behavioural plasticity.

## Discussion

How mechanisms of robust generation of specific neuron types are balanced with the ability to drive plastic changes in gene expression and behaviour and how these mechanisms are related to the evolution of neuron types is not well understood. We have found that in *C. elegans*, robust development of VC4 and VC5 cholinergic neurons requires H3K9me epigenetic signalling to suppress a 5HT reuptake phenotype. H3K9me-mediated repression of undesired effector genes in the differentiation of both neuronal and non-neuronal tissues has been reported in the past^16^. In addition, we show this process is also plastic, as epigenetic repression of VC4 and VC5 5HT reuptake phenotype can be released when worms develop under specific environmental conditions that promote an alternative yet highly robust serotonergic phenotype. These findings demonstrate a neuronal plasticity mechanism regulated through H3K9me epigenetic repression, akin to other plastic responses described in the immune system^50–52^.

Our results in *C. elegans* prompted us to systematically investigate the distribution of the VC4 and VC5 5HT phenotype in the *Caenorhabditis* genus. Under laboratory conditions, we found that the *Angaria* group of species share strong and constitutive 5HT staining in VC4 and VC5 neurons. This robust phenotype is provided by the emergence of an enhancer in the *mod-5*/Sert locus, which is active in VC4 and VC5 neurons, allowing for the robust 5HT reuptake phenotype. When placed in the *C. elegans* genomic context, this enhancer is sufficient to overcome H3K9me repression in VC4 and VC5 neuron development without the requirement of any environmental trigger and is recruited as a direct target of the neuron-type gene regulatory network controlled by UNC-4/UNCX TF. Thus, the emergence of this new regulatory module is sufficient to produce a shift from plastic to a constitutive trait. cis-regulatory changes in the neuropeptide PDF have also been shown recently to underlie the evolution of circadian behaviour plasticity in *Drosophila* species^53^. Our results provide an example of the evolution of neuron-type specific gene expression and neuronal function and describe its underlying gene regulatory mechanism, giving experimental support to previous models^1,2^.

Moreover, we identify behavioural differences that can be directly attributed to this new trait: the ability to block egg-laying induction by external sources of 5HT. We speculate that this neuronal function might be adaptative as tight temporal control of egg-laying and egg retention in harsh environments is important for embryonic viability and population competition^54^. We propose that evolution of this constitutive trait in *Angaria* species could be driven, at least in part, by exposure to high levels of environmental 5HT found in some plants and fruits, which can be further increased upon insect feeding, or fungal infections^33–39^. The specific constitutive trait for *Angaria* species is intriguing as they share some natural habitats with other *Caenorhabditis* species^55^. We speculate that the plastic responses to exogenous 5HT we found for *C. elegans* might be related to its generalist lifestyle and the fact that it is frequently found in slug’s intestines where it can survive intestinal transit^56^. This particular behaviour has been suggested to be a dispersal mechanism. The intestine is the major source of 5HT in the body, including snails and slugs^57,58^. Considering that phenotypic plasticity is favoured in changing environments where predictive cues are present, it is tempting to speculate *C. elegans* VC4 and VC5 serotonergic plasticity might be key to induce egg retention when present in intestinal environments, which will favour offspring survival but not in external habitats that will favour egg deposition, population dispersal and colonisation of new food resources. The gain and subsequent loss of plasticity are proposed to have a role in the emergence of phenotypic novelty, a concept known as genetic assimilation^59^. The lack of a plastic response to exogenous 5HT in *C. briggsae* and *C. tropicalis* suggests that *C. elegans* might have acquired this response. We cannot assess if a plastic state preceded constitutive VC4 and VC5 5HT staining in *Angaria* species. Nevertheless, constitutive serotonergic staining in VC4 and VC5 neurons is present in several non-*Caenorhabditis* nematodes of diverse origin^30^, suggesting this phenotype has been gained or lost repeatedly, further supporting its evolvability. Importantly, we provide specific genetic mechanisms underlying both trait plasticity (facultative epigenetic silencing) and canalisation (enhancer emergence refractory to epigenetic silencing).

Finally, our results also bear biomedical implications. Sert genetic variants and the use of SERT inhibitors during pregnancy have been linked to pathological conditions such as depression, anxiety, autism or pain perception^60,61^. Importantly, there is also a key interplay between Sert genetic variants and environmental modulation (vulnerability or protection to disease)^62^. The mechanisms directing dynamic MOD-5/SERT expression in cortical neurons are unknown, as well as whether environmental stimuli can modulate its repression under physiological or pathological conditions. Indeed, mouse retinal ganglion cells, which transiently express SERT in development, reactivate its expression after optic nerve injury, increasing the vulnerability of these neurons^63^. Importantly, H3K9me pathway has also been genetically linked to psychiatric disorders^64,65^. In light of our results, we speculate that plastic H3K9me epigenetic silencing of Sert locus might be phylogenetically conserved in mammals, could potentially be environmentally modulated and involved in 5HT related neuropsychiatric disorders.

## Acknowledgements

We thank Oscar Marín, Oliver Hobert, Christian Braendle, Santiago Ramón and Santiago Vernia for their comments on the manuscript. James Rand and Janet Duerr for generously sharing the UNC-17 antibody. Laura Chirivella and Ana Roig for technical assistance. Inés Carrera, Isabel del Pino, Eduardo Leyva and Nuria Flames lab members for feedback on the project.

## References

1. Arendt, D. The evolution of cell types in animals: Emerging principles from molecular studies. Nature Reviews Genetics Preprint at 10.1038/nrg2416 (2008).

2. Arendt, D., Bertucci, P. Y., Achim, K. & Musser, J. M. Evolution of neuronal types and families. Curr Opin Neurobiol 56, 144–152 (2019).

3. Gaspar, P., Cases, O. & Maroteaux, L. The developmental role of serotonin: News from mouse molecular genetics. Nature Reviews Neuroscience vol. 4 1002–1012 Preprint at 10.1038/nrn1256 (2003).

4. Wong, F. K. et al. Serotonergic regulation of bipolar cell survival in the developing cerebral cortex. Cell Rep 40, (2022).

5. GarcíFrigola, C. & Herrera, E. Zic2 regulates the expression of Sert to modulate eye-specific refinement at the visual targets. EMBO Journal 29, 3170–3183 (2010).

6. Moroz, L. L., Romanova, D. Y. & Kohn, A. B. Neural versus alternative integrative systems: Molecular insights into origins of neurotransmitters. Philosophical Transactions of the Royal Society B: Biological Sciences 376, (2021).

7. Memar, N. et al. Twenty million years of evolution: The embryogenesis of four Caenorhabditis species are indistinguishable despite extensive genome divergence. Dev Biol 447, 182–199 (2019).

8. Zhao, Z. et al. Comparative analysis of embryonic cell lineage between Caenorhabditis briggsae and Caenorhabditis elegans. Dev Biol 314, 93–99 (2008).

9. Chase, D. L. & Koelle, M. R. Biogenic amine neurotransmitters in C. elegans. WormBook : the online review of C. elegans biology 1–15 Preprint at 10.1895/wormbook.1.132.1 (2007).

10. Curran, K. P. & Chalasani, S. H. Serotonin circuits and anxiety: What can invertebrates teach us? Invertebrate Neuroscience vol. 12 81–92 Preprint at 10.1007/s10158-012-0140-y (2012).

11. Ranganathan, R., et al. Mutations in the Caenorhabditis Elegans Serotonin Reuptake Transporter MOD-5 Reveal Serotonin-Dependent and-Independent Activities of Fluoxetine. The Journal of Neuroscience vol. 21 (2001).

12. Jafari, G., Xie, Y., Kullyev, A., Liang, B. & Sze, J. Y. Regulation of extrasynaptic 5-HT by serotonin reuptake transporter function in 5-HT-absorbing neurons underscores adaptation behavior in Caenorhabditis elegans. Journal of Neuroscience 31, 8948–8957 (2011).

13. Duerr, J. S., Gaskin, J. & Rand, J. B. Identified Neurons in C. Elegans Coexpress Vesicular Transporters for Acetylcholine and Monoamines. http://www.ajpcell.org (2001).

14. Duerr, J. S., et al. The Cat-1 Gene of Caenorhabditis Elegans Encodes a Vesicular Monoamine Transporter Required for Specific Monoamine-Dependent Behaviors. (1998).

15. Poole, R. J., Flames, N. & Cochella, L. Neurogenesis in Caenorhabditis elegans. Genetics (2024) doi:10.1093/genetics/iyae116.

16. Padeken, J., Methot, S. P. & Gasser, S. M. Establishment of H3K9-methylated heterochromatin and its functions in tissue differentiation and maintenance. Nature Reviews Molecular Cell Biology vol. 23 623–640 Preprint at 10.1038/s41580-022-00483-w (2022).

17. Methot, S. P. et al. H3K9me selectively blocks transcription factor activity and ensures differentiated tissue integrity. Nat Cell Biol 23, 1163–1175 (2021).

18. Nicetto, D. et al. H3K9me3-heterochromatin loss at protein-coding genes enables developmental lineage specification. Science (1979) 363, 294–297 (2019).

19. Allan, R. S. et al. An epigenetic silencing pathway controlling T helper 2 cell lineage commitment. Nature 487, 249–253 (2012).

20. Lyons, D. B. et al. Heterochromatin-mediated gene silencing facilitates the diversification of olfactory neurons. Cell Rep 9, 884–892 (2014).

21. Magklara, A. et al. An epigenetic signature for monoallelic olfactory receptor expression. Cell 145, 555–570 (2011).

22. Ahringer, J. & Gasser, S. M. Repressive chromatin in caenorhabditis elegans: Establishment, composition, and function. Genetics 208, 491–511 (2018).

23. Lloret-Fernández, C. et al. A transcription factor collective defines the HSN serotonergic neuron regulatory landscape. Elife 7, (2018).

24. Desai, C., Garriga, G., Mclntire, S. L. & Horvitz, H. R. A genetic pathway for the development of the Caenorhabditis elegans HSN motor neurons. Nature 336, 638–646 (1988).

25. Finney, M. & Ruvkun, G. The unc-86 gene product couples cell lineage and cell identity in C. elegans. Cell 63, 895–905 (1990).

26. Kudron, M. M. et al. The modern resource: genome-wide binding profiles for hundreds of Drosophila and Caenorhabditis elegans transcription factors. Genetics 208, 937–949 (2018).

27. Zheng, C., Karimzadegan, S., Chiang, V. & Chalfie, M. Histone Methylation Restrains the Expression of Subtype-Specific Genes during Terminal Neuronal Differentiation in Caenorhabditis elegans. PLoS Genet 9, e1004017 (2013).

28. Lickteig, K. M. et al. Regulation of neurotransmitter vesicles by the homeodomain protein UNC-4 and its transcriptional corepressor UNC-37/groucho in Caenorhabditis elegans cholinergic motor neurons. Journal of Neuroscience (2001) doi:10.1523/jneurosci.21-06-02001.2001.

29. Bany, I. A., Dong, M.-Q. & Koelle, M. R. Genetic and Cellular Basis for Acetylcholine Inhibition of Caenorhabditis Elegans Egg-Laying Behavior. (2003).

30. Loer, C. M. & Rivard, L. Evolution of neuronal patterning in free-living rhabditid nematodes I: Sex-specific serotonin-containing neurons. Journal of Comparative Neurology (2007) doi:10.1002/cne.21288.

31. Schafer, W. R. Egg-laying. WormBook : the online review of C. elegans biology 1–7 Preprint at 10.1895/wormbook.1.38.1 (2005).

32. Schulenburg, H. & Félix, M. A. The natural biotic environment of Caenorhabditis elegans. Genetics 206, 55–86 (2017).

33. Zhang, H. et al. A specific primed immune response in red palm weevil, Rhynchophorus ferrugineus, is mediated by hemocyte differentiation and phagocytosis. Dev Comp Immunol 131, (2022).

34. Erland, L. A. E., Turi, C. E. & Saxena, P. K. Serotonin: An ancient molecule and an important regulator of plant processes. Biotechnology Advances vol. 34 1347–1361 Preprint at 10.1016/j.biotechadv.2016.10.002 (2016).

35. Ramakrishna, A., Giridhar, P. & Ravishankar, G. A. Phytoserotonin: A review. Plant Signaling and Behavior vol. 6 800–809 Preprint at 10.4161/psb.6.6.15242 (2011).

36. Dangol, A. et al. Characterizing serotonin biosynthesis in Setaria viridis leaves and its effect on aphids. Plant Mol Biol 109, 533–549 (2022).

37. Lu, H. P. et al. Resistance of rice to insect pests mediated by suppression of serotonin biosynthesis. Nat Plants 4, 338–344 (2018).

38. Shmukler, Y. B. & Nikishin, D. A. Non-Neuronal Transmitter Systems in Bacteria, Non-Nervous Eukaryotes, and Invertebrate Embryos. Biomolecules vol. 12 Preprint at 10.3390/biom12020271 (2022).

39. Saremba, B. M. et al. Plant signals during beetle (Scolytus multistriatus) feeding in American elm (Ulmus americana Planch). Plant Signal Behav 12, (2017).

40. Gilbert, S. F., Bosch, T. C. G. & Ledón-Rettig, C. Eco-Evo-Devo: Developmental symbiosis and developmental plasticity as evolutionary agents. Nature Reviews Genetics vol. 16 611–622 Preprint at 10.1038/nrg3982 (2015).

41. Sudhaus, W., Kiontke, K. & Giblin-Davis, R. M. Description of Caenorhabditis angaria n. sp. (Nematoda: Rhabditidae), an associate of sugarcane and palm weevils (Coleoptera: Curculionidae). Nematology 13, 61–78 (2011).

42. Ramakrishna, A., Giridhar, P. & Ravishankar, G. A. Phytoserotonin: A review. Plant Signaling and Behavior vol. 6 800–809 Preprint at 10.4161/psb.6.6.15242 (2011).

43. O’Donnell, M. P., Fox, B. W., Chao, P. H., Schroeder, F. C. & Sengupta, P. A neurotransmitter produced by gut bacteria modulates host sensory behaviour. Nature 583, 415–420 (2020).

44. Horvitz, H. R., Chalfie, M., Trent, C., Sulston, J. E. & Evans, P. D. Serotonin and Octopamine in the Nematode *Caenorhabditis elegans*. Science (1979) 216, 1012–1014 (1982).

45. Hobson, R. J. et al. SER-7, a Caenorhabditis elegans 5-HT7-like receptor, is essential for the 5-HT stimulation of pharyngeal pumping and egg laying. Genetics 172, 159–169 (2006).

46. Shyn, S. I., Kerr, R. & Schafer, W. R. Serotonin and Go Modulate Functional States of Neurons and Muscles Controlling C. elegans Egg-Laying Behavior. Current Biology 13, 1910–1915 (2003).

47. Carnell, L., Illi, J., Hong, S. W. & McIntire, S. L. The G-protein-coupled serotonin receptor SER-1 regulates egg laying and male mating behaviors in Caenorhabditis elegans. Journal of Neuroscience 25, 10671–10681 (2005).

48. Flames, N. & Hobert, O. Transcriptional Control of the Terminal Fate of Monoaminergic Neurons. Annu Rev Neurosci (2011) doi:10.1146/annurev-neuro-061010-113824.

49. Toker, I. A. et al. Molecular patterns of evolutionary changes throughout the whole nervous system of multiple nematode species. Preprint at 10.1101/2024.11.23.624988 (2024).

50. van Essen, D., Zhu, Y. & Saccani, S. A Feed-Forward Circuit Controlling Inducible NF-κB Target Gene Activation by Promoter Histone Demethylation. Mol Cell 39, 750–760 (2010).

51. Xiao, X. et al. The Costimulatory Receptor OX40 Inhibits Interleukin-17 Expression through Activation of Repressive Chromatin Remodeling Pathways. Immunity 44, 1271–1283 (2016).

52. Hachiya, R. et al. The H3K9 methyltransferase Setdb1 regulates TLR4-mediated inflammatory responses in macrophages. Sci Rep 6, (2016).

53. Shahandeh, M. P. et al. Circadian plasticity evolves through regulatory changes in a neuropeptide gene. Nature (2024) doi:10.1038/s41586-024-08056-x.

54. Mignerot, L. et al. Natural variation in the Caenorhabditis elegans egg-laying circuit modulates an intergenerational fitness trade-off. Elife 12, (2023).

55. Sloat, S. A. et al. Caenorhabditis nematodes colonize ephemeral resource patches in neotropical forests. Ecol Evol 12, (2022).

56. Petersen, C. et al. Travelling at a slug’s pace: Possible invertebrate vectors of Caenorhabditis nematodes. BMC Ecol 15, (2015).

57. Á S Z L Ó H E R N Á D I, L., O S E R D É Ly I, L. A. J. & Ly E L E K, R. O. The Organization of Serotonin-, Dopamine-, and FMRFamide-Containing Neuronal Elements and Their Possible Role in the Regulation of Spontaneous Contraction of the Gastrointestinal Tract in the Snail Helix Pomatia Monoamines and FMRFamide in the Gastropod Intestine HERNÁDI, ERDÉLYI, HIRIPI, and ELEKES. (1998).

58. Rao, M. & Gershon, M. D. Enteric nervous system development: what could possibly go wrong? Nature Reviews Neuroscience vol. 19 552–565 Preprint at 10.1038/s41583-018-0041-0 (2018).

59. Waddington, C. H. Genetic Assimilation of an Acquired Character. Source: Evolution vol. 7 (1953).

60. Murphy, D. L. & Moya, P. R. Human serotonin transporter gene (SLC6A4) variants: Their contributions to understanding pharmacogenomic and other functional G × G and G × e differences in health and disease. Current Opinion in Pharmacology vol. 11 3–10 Preprint at 10.1016/j.coph.2011.02.008 (2011).

61. Murphy, D. L. & Lesch, K. P. Targeting the murine serotonin transporter: Insights into human neurobiology. Nature Reviews Neuroscience vol. 9 85–96 Preprint at 10.1038/nrn2284 (2008).

62. Hankin, B. L. et al. Differential susceptibility in youth: Evidence that 5-HTTLPR x positive parenting is associated with positive affect for better and worse. Transl Psychiatry 1, (2011).

63. Kingston, R., Amin, D., Misra, S., Gross, J. M. & Kuwajima, T. Serotonin transporter-mediated molecular axis regulates regional retinal ganglion cell vulnerability and axon regeneration after nerve injury. PLoS Genet 17, (2021).

64. Walker, D. M., Cates, H. M., Heller, E. A. & Nestler, E. J. Regulation of chromatin states by drugs of abuse. Current Opinion in Neurobiology vol. 30 112–121 Preprint at 10.1016/j.conb.2014.11.002 (2015).

65. Peña, C. J., Bagot, R. C., Labonté, B. & Nestler, E. J. Epigenetic signaling in psychiatric disorders. Journal of Molecular Biology vol. 426 3389–3412 Preprint at 10.1016/j.jmb.2014.03.016 (2014).

