## Supplementary Figures for "Gene regulatory mechanisms underlying evolutionary adaptations of homologous neuronal cell types"

### Extended Data Figures

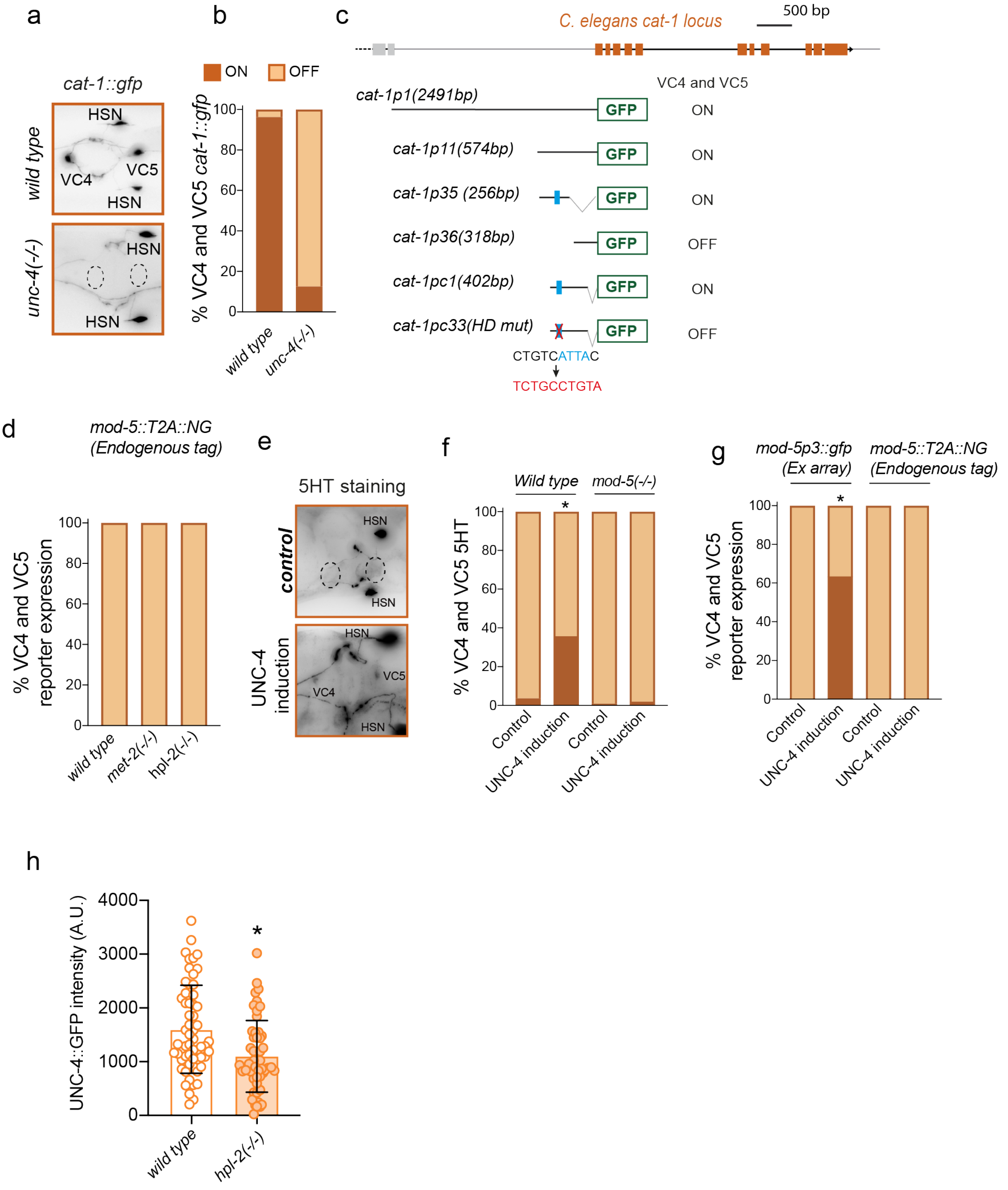

**Extended Data Figure 1. UNC-4 role in VC4 and VC5 serotonergic phenotype. a**) Micrographs showing *cat-1::gfp* reporter expression in *wildtype* VC4 and VC5 neurons and loss of expression in *unc-4(gk705)* mutant background. **b)** Quantification of (a). **c)** *cat-1* VC4 and VC5 minimal CRM (*cat-1pc1*) contains a functional HD binding site. HD site is depicted in blue, modified sequence is shown in red **d)** Quantification of *mod-5::T2A::NeonGreen* expression. *met-2(n4256)* and *hpl-2(tm1489)* mutans show no detectable expression in VC4 and VC5 neurons. **e)** Micrograph of *C. elegans* control animals and animals with overexpression of UNC-4 which induces VC4 and VC5 serotonergic phenotype. **f)** Quantification of VC4 and VC5 serotonergic staining upon UNC-4 induction in *wild type* and *mod-5(knu383)* mutant animals. UNC-4 overexpression drives VC4 and VC5 serotonergic phenotype in a *mod-5* dependent manner. **g)** Quantification of *Cel_mod-5prom3::gfp* multicopy extrachromosomal array reporter and endogenously tagged *mod-5::T2A::NeonGreen* expression upon UNC-4 induction. **h)** Quantification of endogenously tagged UNC-4::GFP intensity in VC4 and VC5 neurons*. hpl-2(tm1489)* mutants show a significant decrease in GFP intensity compared to controls.

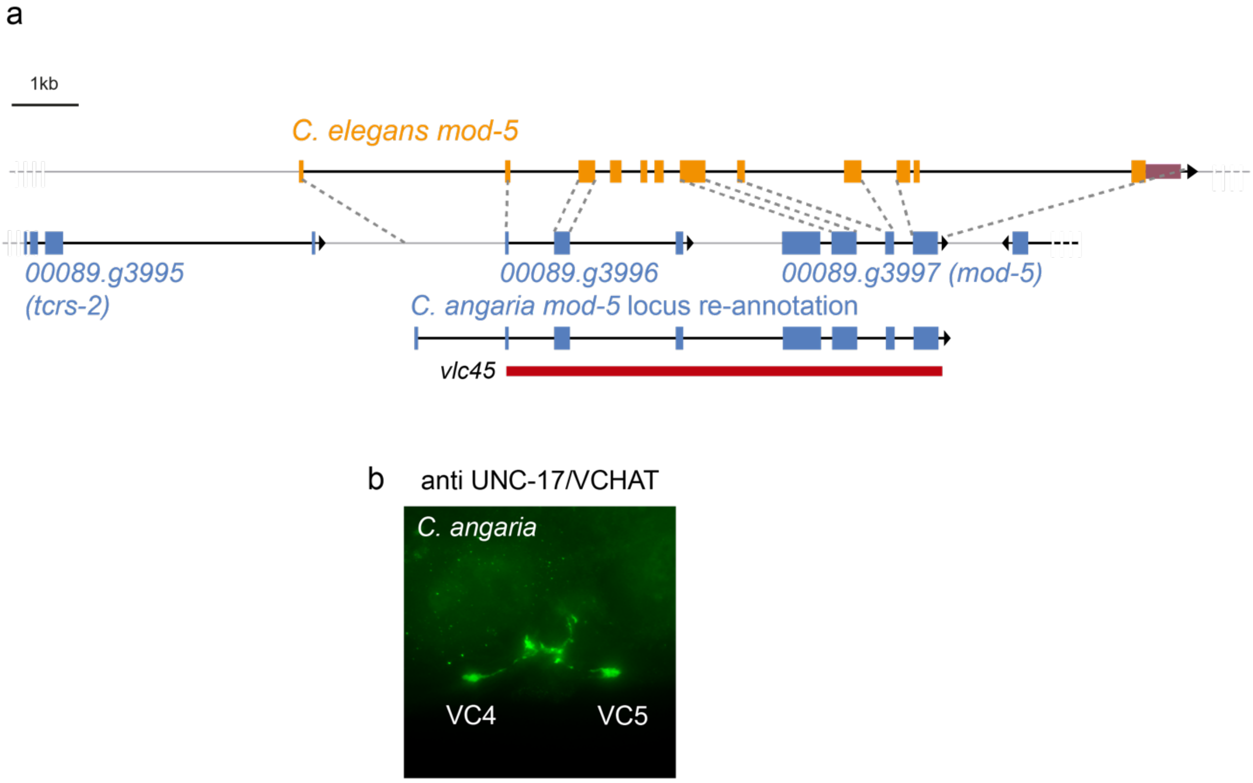


**Extended Data Figure 2. *C. angaria mod-5* locus re-annotation and characterization of *C. angaria* VC4 and VC5 neurons cholinergic phenotype. a**) Schematic of *C. angaria* genomic region including *mod-5* gene (blue) and alignment to *C. elegans mod-5* locus (orange). Original *C. angaria mod-5* gene was annotated as gene 0089.g3997. Alignment with *C. elegans* shows that upstream gene 0089.g3996 contains homology to additional *mod-5* exons. Finally, homology to *Cel* Exon1 is also found in *C. angaria* genome sequence. Proposed re-annotation of the locus is depicted below. *Cang mod-5(vlc45)* is a deletion allele from exon2 until the end of the locus, generating a null mutant. *PRJNA51225 C. angaria* genome from Wormbase Parasite, contig *Cang_2012_01_13_00089*. **b**) Immunostaining of *C. angaria* animals with anti-UNC-17/VCHAT antibody showing VC4 and VC5 express also cholinergic markers.

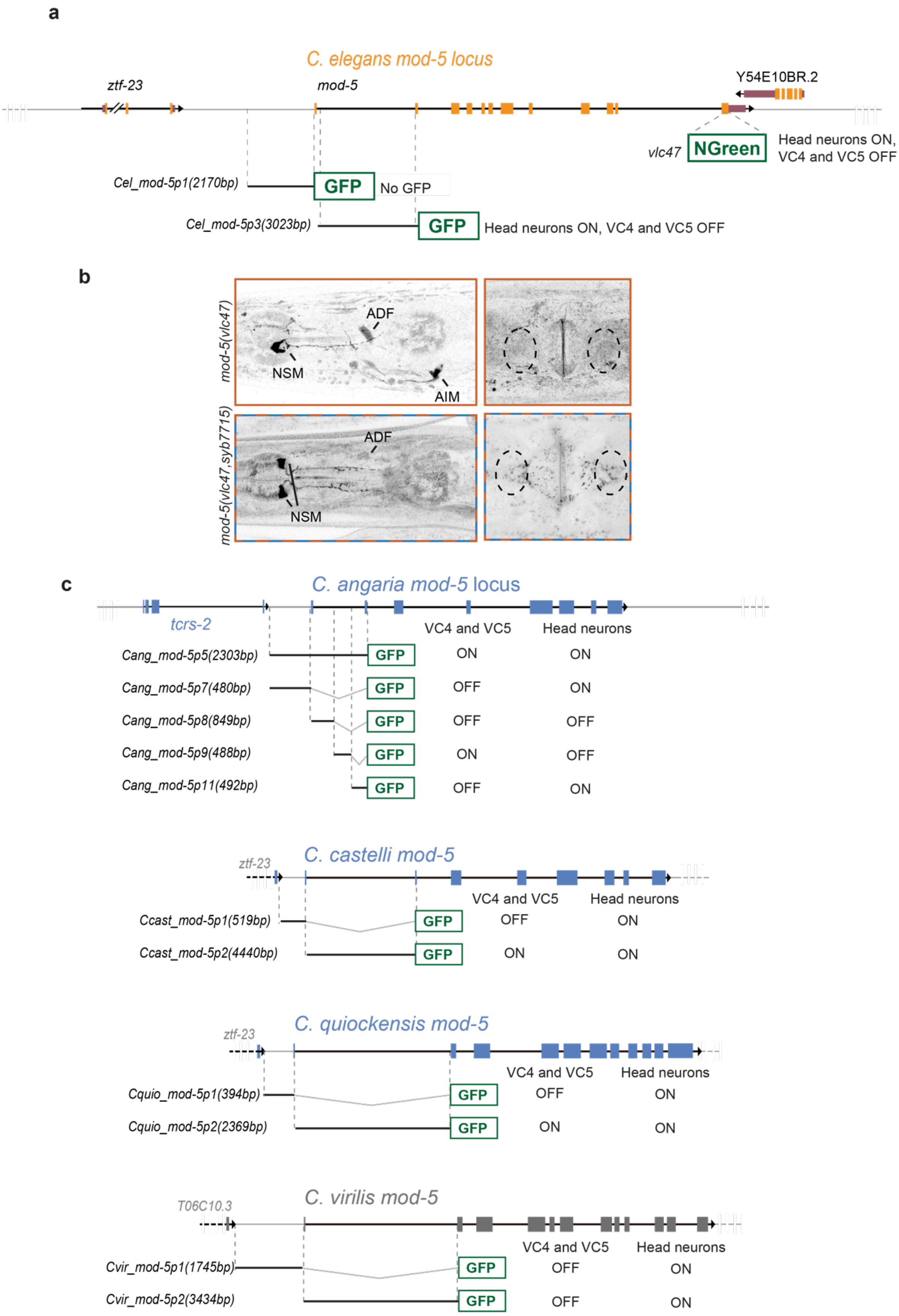

a
C. elegans mod-5 locus
ztf-23
mod-5
Y54E10BR.2
vlc47
NGreen
Head neurons ON,
VC4 and VC5 OFF
Cel_mod-5p1(2170bp)
GFP
No GFP
Cel_mod-5p3(3023bp)
GFP
Head neurons ON, VC4 and VC5 OFF
b
mod-5(vlc47)
ADF
NSM
AIM
mod-5(vlc47,syb7715)
ADF
NSM
c
C. angaria mod-5 locus
tcrs-2
VC4 and VC5
Head neurons
Cang_mod-5p5(2303bp)
GFP
ON
ON
Cang_mod-5p7(480bp)
GFP
OFF
ON
Cang_mod-5p8(849bp)
GFP
OFF
OFF
Cang_mod-5p9(488bp)
GFP
ON
OFF
Cang_mod-5p11(492bp)
GFP
OFF
ON
C. castelli mod-5
ztf-23
VC4 and VC5
Head neurons
Ccast_mod-5p1(519bp)
GFP
OFF
ON
Ccast_mod-5p2(4440bp)
GFP
ON
ON
C. quiockensis mod-5
ztf-23
VC4 and VC5
Head neurons
Cquio_mod-5p1(394bp)
GFP
OFF
ON
Cquio_mod-5p2(2369bp)
GFP
ON
ON
C. virilis mod-5
T06C10.3
VC4 and VC5
Head neurons
Cvir_mod-5p1(1745bp)
GFP
OFF
ON
Cvir_mod-5p2(3434bp)
GFP
OFF
ON

**Extended Data Figure 3. *mod-5* CRM reporter constructs from *Caenorhabditis* species and their activity in *C. elegans* transgenic animals. a**) *C. elegans mod-5* locus, analysed reporters and summary of expression. ***b**) C. elegans mod-5(vlc47)* endogenously tagged allele drives detectable NeonGreen expression in ADF, NSM, AIM and RIH serotonergic neurons similar to previous reports (Jafari et al., 2011). *C. elegans mod-5(vlc47 syb7715)* strain*,* in which *mod-5* first intron is substituted by *C. angaria* intron 1 sequence shows strong NSM expression, decreased ADF and AIM expression and undetectable VC4 and VC5 expression despite strong 5HTergic immunostaining. **c**) *mod-5* locus in *Angaria* species (blue) and non-*angaria* species (grey) and analysed constructs with summary of expression. *mod-5* first intron contains enhancers active in head neurons in all species, while VC4 and VC5 activity is only observed in *Angaria* species reporters. See Supplementary information for primary data.

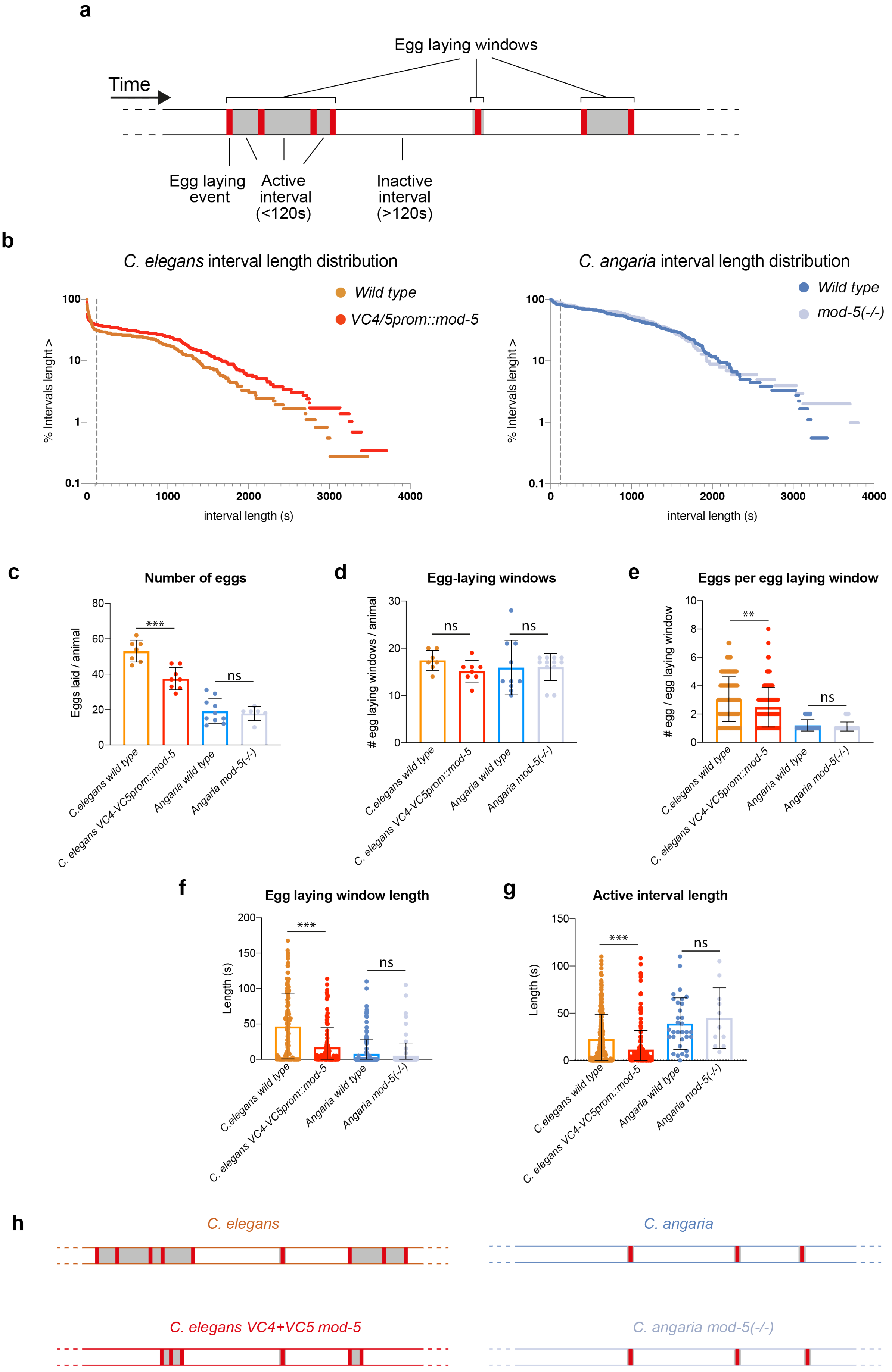
a
Egg laying windows
Time
Egg laying event
Active interval (<120s)
Inactive interval (>120s)
b
C. elegans interval length distribution
Wild type
VC4/5prom::mod-5
% Intervals lenght >
interval length (s)
C. angaria interval length distribution
Wild type
mod-5(-/-)
c
Number of eggs
Eggs laid / animal
***
ns
C.elegans wild type
C. elegans VC4-VC5prom::mod-5
Angaria wild type
Angaria mod-5(-/-)
d
Egg-laying windows
### egg laying windows / animal
ns
e
Eggs per egg laying window
### egg / egg laying window
**
f
Egg laying window length
Length (s)
g
Active interval length
h
C. elegans
C. angaria
C. elegans VC4+VC5 mod-5
C. angaria mod-5(-/-)

**Extended Data Figure 4. Egg laying dynamics in *C. elegans* and *C. angaria. a)*** Schematic representation of egg laying behaviour and terms used in the analysis. Animals were recorded moving freely in OP50 plates for 6h and egg laying events (shown as red lines) were manually annotated. *C. elegans* egg-laying behaviour transitions from active periods of egg laying (shown in grey) to inactive periods with no egg laying. Inactive intervals are assigned when the time between two egg-laying events is longer than two minutes. An egg-laying window is defined by the time comprised between the first and the last egg-laying event with intervals shorter than two minutes (active intervals). **b)** Interval length distribution in *C. elegans* wild type compared to transgenic animals expressing *mod-5* cDNA in VC neurons (left graph) or *C. angaria* wild type (RGD1) compared to *C. angaria mod-5(vlc45)* mutant (right graph). Distributions are different between species but not when comparing genotypes within the same species. Dotted lines marks 120s intervals, dividing active from inactive intervals. **c)** Quantification of total number of laid eggs. *C. elegans* transgenic animals expressing *mod-5* cDNA in VC neurons lay significantly less eggs than *C. elegans* wild type T-test was used). **d**) There are no significant differences in the number of egg laying windows among groups (T-test was used). **e)** Quantification of number of eggs laid per Egg laying window shows significantly smaller number of eggs laid by *C. elegans* transgenic animals expressing *mod-5* cDNA in VC neurons compared to wild type T-test was used. **f**) Egg laying window length quantification, showing significantly shorter windows in *C. elegans* transgenic animals compared to wild type Wilcoxon test was used. **g**) Active interval length is also significantly shorter in *C. elegans* transgenic animals compared to wild type Wilcoxon test was used. **h**) Summary of egg laying behaviours in the different species and genetic backgrounds, *C. elegans* transgenic animals show shorter active intervals, shorter egg laying windows and less egg laying events compared to wild type, however, no differences are found between *C. angaria* and *C. angaria mod-5(vlc45)* mutant egg laying behaviour under this laboratory conditions. See Supplementary information for primary data.

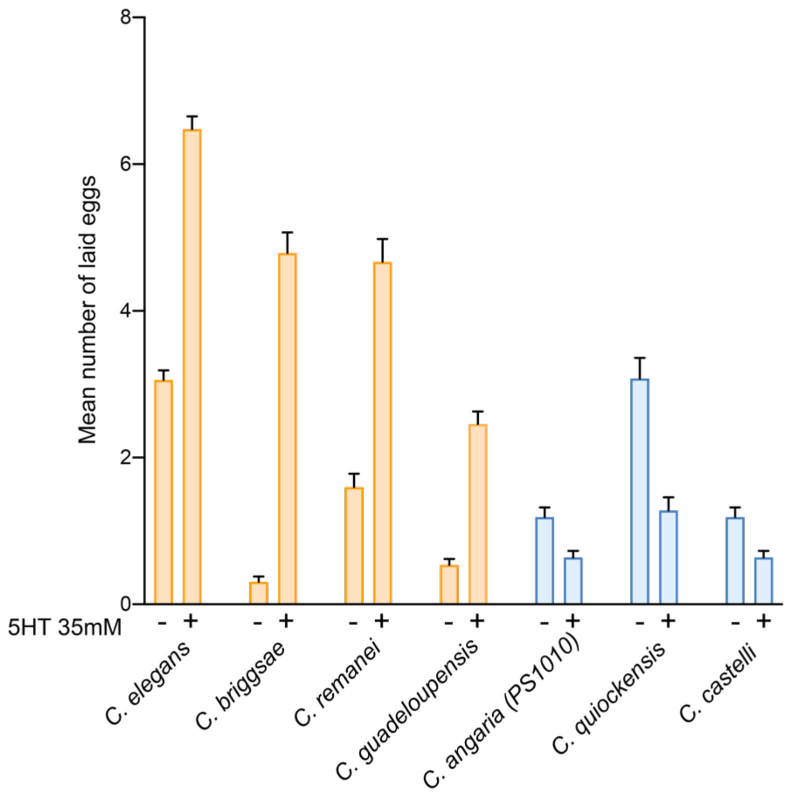


**Extended Data Figure 5. Egg laying quantification upon acute 5HT exposure in different *Caenorhabditis* species.** Worms were placed in M9 or M9 with 35 mM for 60 minutes and then laid eggs were quantified. Mean and SD number of eggs from 3 different experiments is represented. There is an anti-correlation between VC4 and VC5 serotonergic phenotype and egg-laying induction by external serotonin, *Angaria* species (in blue) show no induction or inhibition of egg-laying upon external serotonin exposure while *non-Angaria* species (orange) show strong egg-laying induction. See Supplementary information for primary data.

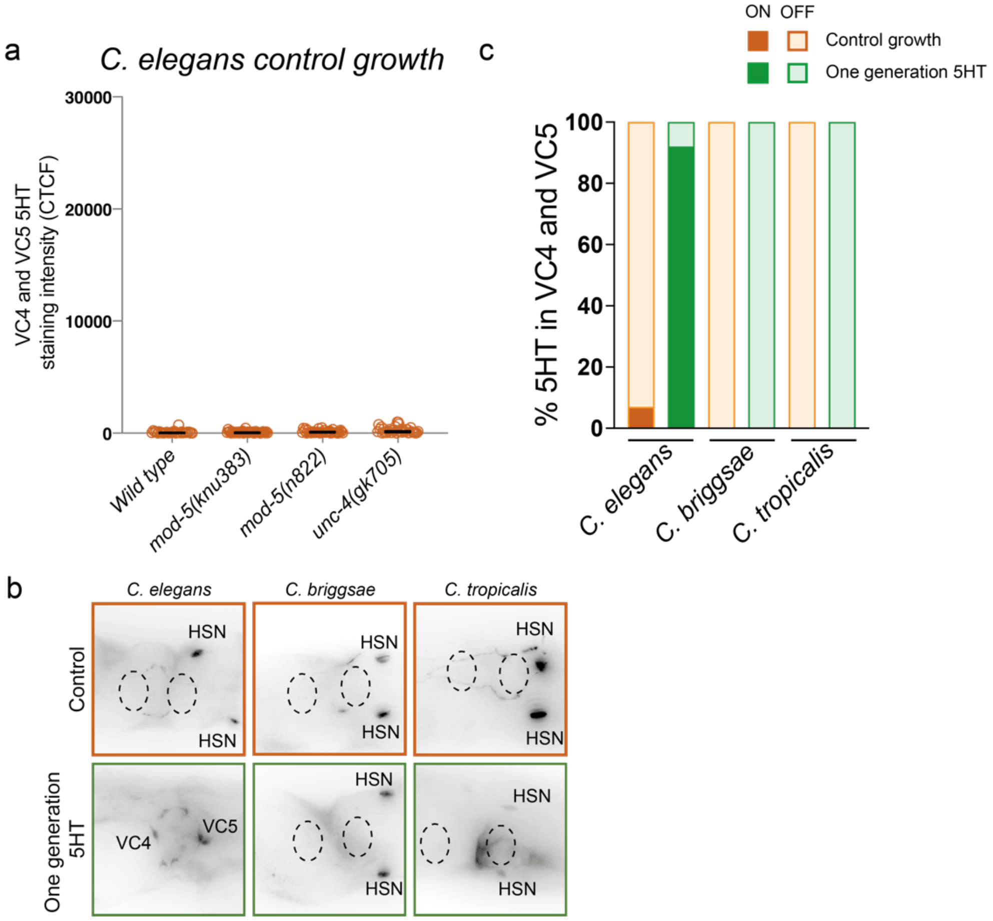


**Extended Data Figure 6. Study of VC4 and VC5 serotonergic plasticity in different Caenorhabditis species. a)** wild type, two *mod-5* alleles and *unc-4* mutants quantification of VC4 and VC5 5HT staining intensity under control growth condition showing no detectable levels of 5HT staining. **b)** Representative micrographs of VC4 and VC5 5HT staining in *C. elegans, C. briggsae* and *C. tropicalis* comparing control and growth in high environmental 5HT levels. **c**) Quantification of percentage of animals with detectable 5HT staining in VC4 and VC5. *C. elegans,* but not *C. briggsae* or *C. tropicalis* can acquire the serotonergic phenotype under these conditions
